# SMARCA2 and SMARCA4 participate in DNA damage repair

**DOI:** 10.1101/2024.03.28.587286

**Authors:** Lily Yu, Duo Wu

## Abstract

SMARCA2 and SMARCA4 (SMARCA2/4) are paralogs and act as the key enzymatic subunits in the SWI/SNF complex for chromatin remodeling. However, the role of SMARCA2/4 in DNA damage response remains unclear. Here, we show that SMARCA2/4 relocate to DNA lesions in response to DNA damage, which requires their ATPase activities. Moreover, these ATPase activities are also required for the relocation of other subunits in the SWI/SNF complex to DNA lesions. Interestingly, the relocation of SMARCA2/4 is independent of γH2AX, ATM, ATR, p300/CBP, or PARP1/2, indicating that it may directly recognize DNA lesions as a DNA damage sensor. Lacking SMARCA2/4 prolongs the retention of γH2AX, RNF8 and BRCA1 at DNA lesions and impairs RAD51-dependent homologous recombination repair. Moreover, the treatment of SMARCA2/4 inhibitor sensitizes tumor cells to PARP inhibitor treatment. Collectively, this study reveals SMARCA2/4 as a DNA damage repair factor for double-strand break repair.

## Introduction

Genomic DNA constantly encounters genotoxic stresses that induce DNA damage. In response to DNA damage, the surveillance system of cells is quickly activated(1). DNA lesions are detected by DNA damage sensors, associated with chromatin relaxation for DNA damage repair(2). Several important chromatin remodeling complexes have been reported to participate in DNA damage response. One of them is the SWI/SNF complex(3).

The SWI/SNF complex is evolutionarily conserved in eukaryotes, and plays a key role to regulate the position of nucleosomes on genomic DNA(3–6). This complex is an ATP-dependent chromatin remodeling complex with multiple subunits. The key subunit in the SWI/SNF complex is SMARCA2, an ATPase with the DEGH helicase fold(7,8). It hydrolyzes ATP and provides the energy for the whole complex to disrupt the interaction between histones and DNA, thereby removing nucleosome barriers and exposing naked DNA for numerous biological activities, such as transcription and replication(9,10). SMARCA4 is a paralog of SMARCA2, which contains similar domain architecture and has a redundant function of SMARCA2(2,11–14). In addition to SMARCA2/4, the SWI/SNF complex also has up to 28 other subunits dependent on different biological contexts(7). However, nine of them are known as core subunits that are involved in all the known functions of the SWI/SNF complex(15).

It has been shown that the SWI/SNF complex participates in DNA damage repair. The complex is recruited to DNA double-strand breaks (DSBs) and participates in both non-homologous end joining and homologous recombination repair via sliding nucleosome barriers and exposing naked DNA ends to facilitate the loading of repair machinery(5,6). However, the recruitment of the SWI/SNF complex seems unclear. It has been reported that poly(ADP-ribosyl)ation (aka PARylation) signals or histone acetylation events mediate the recruitment of the SWI/SNF complex to DNA lesions(16–19). However, PARylation is mainly induced by DNA single-strand breaks(20), whereas the SWI/SNF complex is known to participate in DSB repair(5,6). It is also unclear if any specific histone acetylation events are significantly enriched at DNA lesions. Moreover, earlier studies have shown that loss of SMARCA2 is sufficient for impairing DSB repair(5,6). However, it was recently identified that SMARCA4, the paralog of SMARCA2, plays a redundant role in the SWI/SNF complex(21). Since both SMARCA2 and SMARCA4 are ATPase in the SWI/SNF complex, only lacking both of them, but not either of them, can suppress the function of the SWI/SNF complex(21,22). With these unsolved questions, we have carefully examined the recruitment of SMARCA2 and SMARCA4 to DNA lesions as well as their functions in HR repair. We demonstrate that lacking both SMARCA2 and SMARCA4 inhibits HR repair. Moreover, the recruitment of SMARCA2 and SMARCA4 to DNA lesions is independent of PARylation and histone acetylation, but requires the ATPase activities of SMARCA2 and SMARCA4.

## Results

### The enzymatic activities of SMARCA2/4 are essential for the relocation of SMARCA2/4 to DNA lesions

To study the molecular mechanism by which the SWI/SNF complex is recruited to DNA lesions, we examined SMARCA2/4, the key enzymatic subunits in the SWI/SNF complex. Using laser microirradiation assay, we found that both SMARCA2/4 relocated to DNA lesions (Figure 1A).

**Figure 1.**
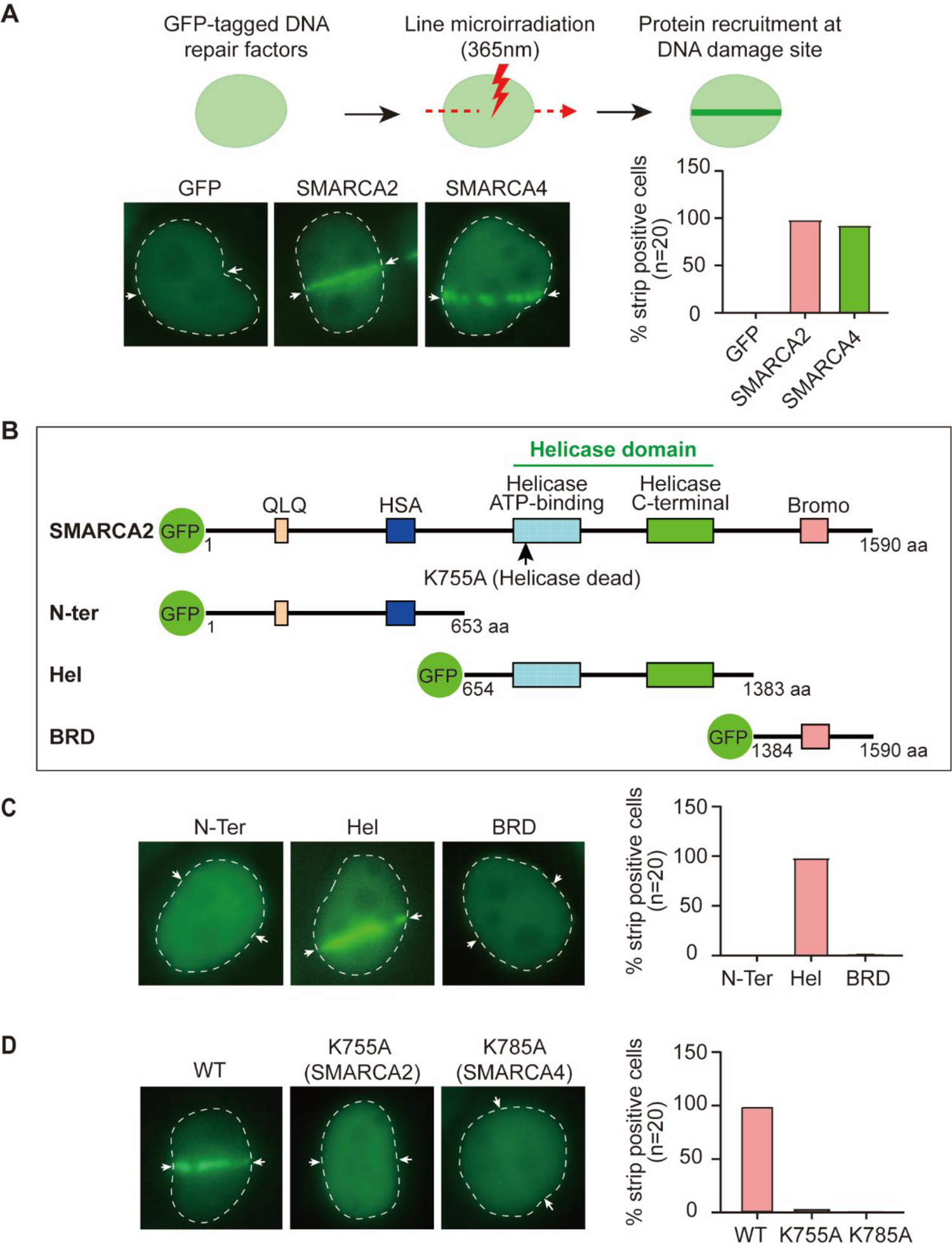
The relocation of SMARCA2/4 to DNA lesions. **(A)** Following laser microirradiation, the laser stripe localization of GFP-SMARCA2/4 was examined. Statistical analyses were included in the lower right panel. **(B)** Schematic of the deletion mutants of SMARCA2. **(C)** The relocation of deletion mutants of SMARCA2 in laser irradiation assays. Statistical analyses were included in the right panel. **(D)** Point mutations that abolished enzymatic activities of SMARCA2. The relocations of these enzymatically dead mutants to DNA lesions were examined.

SMARCA2/4 have several distinct domains, including the N-terminal QLQ motif and HAS domain, which mediate the interaction with other subunits of the SWI/SNF complex; the ATPase/helicase domain in the middle, which is the solo enzymatic domain in the whole complex; the C-terminal bromodomain that is known to recognize acetylated histones (Figure 1B). Since SMARCA2/4 shares identical domain architecture, we generated deletion mutants of SMARCA2, and found that only the ATPase/helicase domain, but not other domains, was able to relocate to DNA lesions (Figure 1C). Moreover, we mutated the key lysine residues in ATPase/helicase domains of SMARCA2/4 to abolish their enzymatic activities, and found that these ATPase activities were required to target SMARCA2/4 to DNA lesions (Figure 1D).

### Relocation of the SWI/SNF complex to DNA lesions is dependent on the ATPase activities of SMARCA2/4

Next, we examined other core subunits of the SWI/SNF complex, including SMARCC1 and SMARCD1. Similar to SMARCA2/4, both SMARCC1 and SMARCD1 relocated to DNA lesions when cells were treated with laser microirradiation (Figure 2A). Interestingly, when we pre-treated cells with FHD286, a dual inhibitor to kill the enzymatic activity of SMARCA2/4, SMARCC1 and SMARCD1 were no longer recruited to DNA lesions in the laser microirradiation assays (Figure 2B), suggesting that the ATPase activity of SMARCA2/4 is also required for targeting other subunits of the SWI/SNF complex to the sites of DNA damage.

**Figure 2.**
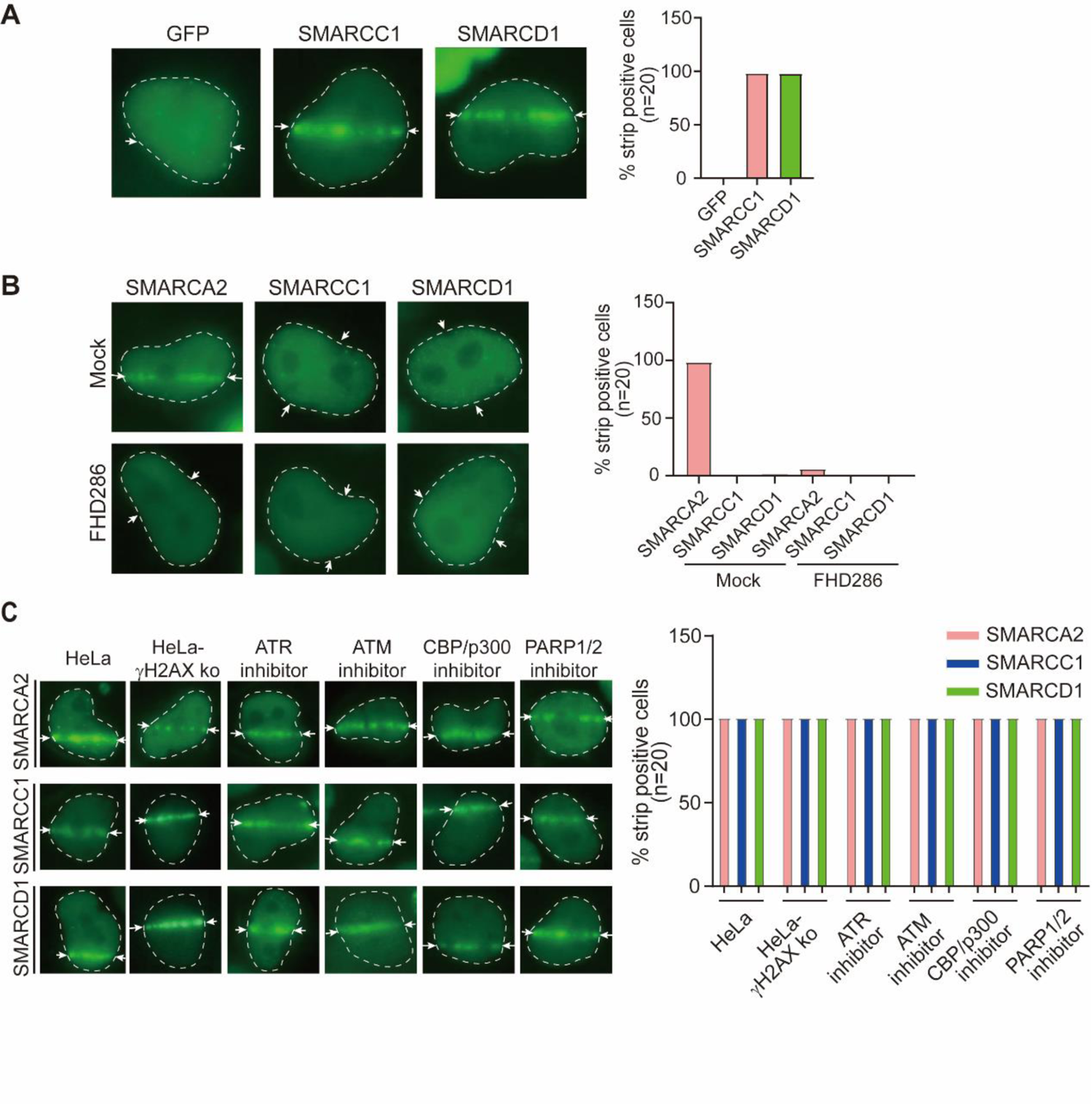
The relocation of the SWI/SNF to DNA lesions. **(A)** The relocation of SMARCC1 and SMARCD1 at laser strip. **(B)** HeLa cells were pre-treated with or without 1 μM FHD286 for two hours. Laser microirradiation was performed to examine the localization of SMARCA2, SMARCC1 and SMARCD1. **(C)** HeLa cells were treated with 1 μM ATMi (ku55933), ATRi (VE-821), CBP/p300i (CBP/p300-IN-21) or PARPi (Olaparib) for two hours followed by laser microirradiation, the localization of SMARCA2, SMARCC1 and SMARCD1 were examined. The localization of the subunits of the SWI/SNF complex was also examined in the H2AX KO HeLa cells.

It has been reported that several DNA repair factors may mediate the recruitment of the SWI/SNF complex to DNA lesions, such as γH2AX, p300/CBP and PARylation(15,19,23). Here, we examined H2AX-deficient cells as well as cells treated with inhibitors of ATM, ATR, p300/CBP, or PARP1/2. To our surprise, the SWI/SNF complex was still recruited to DNA lesions under these conditions (Figure 2C). Only suppression of the ATPase/Helicase domain of SMARCA2/4 abolishes the recruitment of the SWI/SNF complex.

### The enzymatic activity of SMARCA2/4 suppresses the dissociation of DNA damage response factors from DSBs

Earlier studies have shown that the enrichment of DNA damage response factors, such γH2AX, requires SMARCA2(18,19). However, different results have shown that the SWI/SNF complex is the downstream effector of γH2AX at the beginning of the DSB response(24). To further elucidate the role of the SWI/SNF complex in DNA damage response, we used sgRNA to knockout either SMARCA2 or SMARCA4, and treated cells with ionizing radiation (IR) to induce DSBs. However, without either SMARCA2 or SMARCA4, the enrichment of γH2AX at DSBs was not affected (Supplemental Figure S1). Since lacking both SMARCA2 and SMARCA4 causes synthetic lethality for cells, we cannot use sgRNA to obtain SMARCA2/4 DKO cells. Thus, we had to pre-treated cells with FHD286, the potent inhibitor of SMARCA2/4, to transiently suppress the enzymatic activity of SMARCA2/4 before the induction of DSBs. Interestingly, the foci of γH2AX were not suppressed by the treatment of FHD286. Instead, we found the foci of γH2AX existed for a prolonged period following IR treatment (Figure 3). We also examined the downstream factors of γH2AX, including RNF8 and BRCA1. Again, both foci of RNF8 and BRCA1 stayed for an extended period (Figure 3). Moreover, these results were further validated when we treated cells with another potent SMARCA2/4 dual inhibitor Compound 14 (Figure 3). Collectively, these results demonstrate that the suppression of the enzymatic activity of SMARCA2/4 impairs the dissociation of some DNA damage response factors from DSBs.

**Figure 3.**
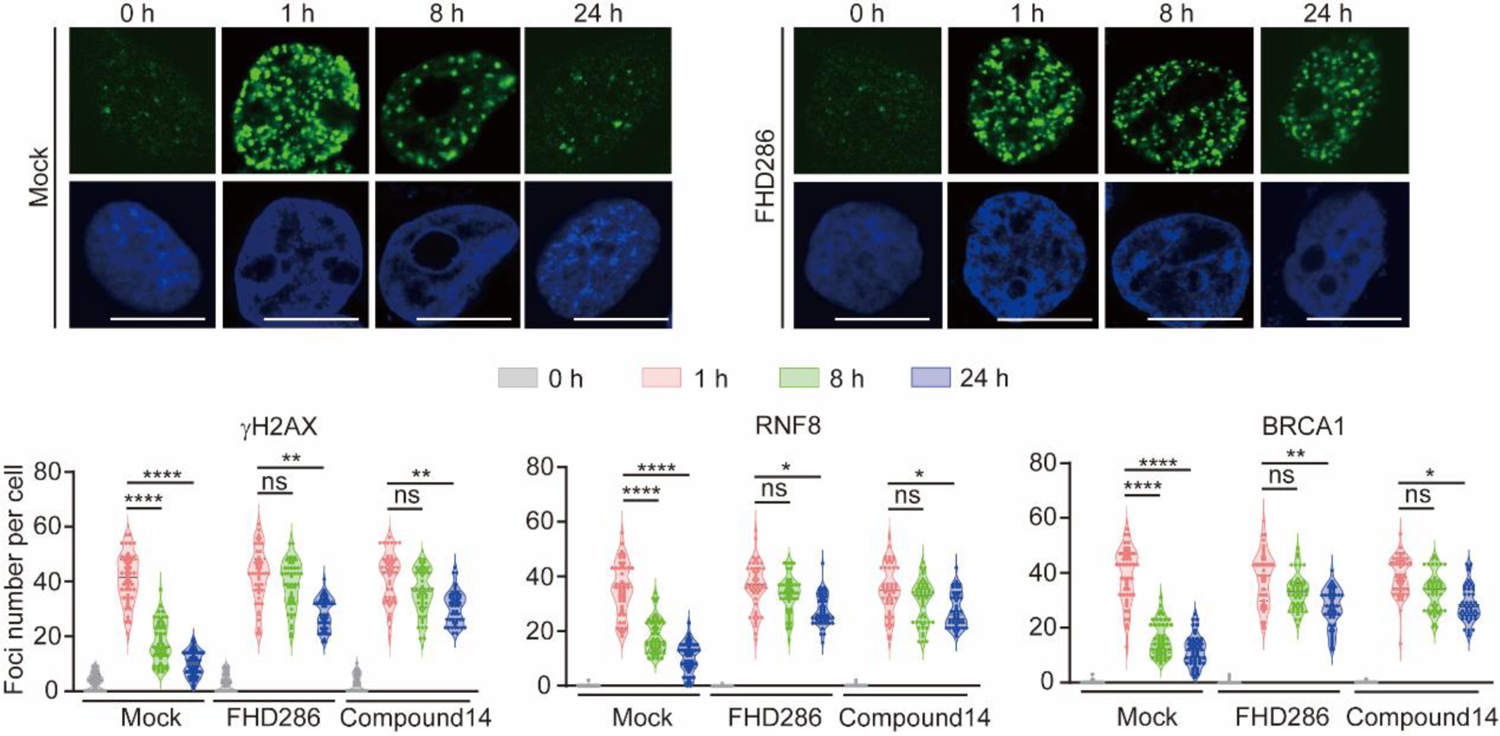
Lacking the enzymatic activities of SMARCA2/4 extends the accumulation of γH2AX, RNF8 and BRCA1. HeLa cells were pre-treated with or without FHD286 (1 μM) or Compound 14 (1 μM) for two hours followed by 5 Gy of IR. Following the indicated recovery time, the foci of γH2AX, RNF8 and BRCA1 were examined by IF. Statistical analyses were shown in the low panels.

### The enzymatic activity of SMARCA2/4 impairs RAD51-mediated HR repair

The prolonged accumulation of DNA damage response factors, such as γH2AX, indicates the defects of DSB repair. It has been shown that the SWI/SNF complex is involved in HR repair. Since SMARCA2/4 are key enzymatic subunits in the SWI/SNF complex, we ask if the ATPase activity of SMARCA2/4 is required for HR repair. During HR repair, RAD51 recognizes ssDNA overhang at DSB ends and motorizes ssDNA overhang to invade into sister chromatids. Thus, RAD51 plays a key role during HR repair. Interestingly, when we treated cells with either FHD286 or Compound 14 to suppress both SMARCA2 and SMARCA4, we found that the IR-induced foci of RAD51 were suppressed (Figure 4A). However, lacking either SMARCA2 or SMARCA4 did not affect the foci formation of RAD51 (Figure 4B). Moreover, we performed a well-established GFP reporter assay to measure the efficacy of HR. Following the treatment of FHD286 or Compound 14, the HR repair was impaired (Figure 4C).

**Figure 4.**
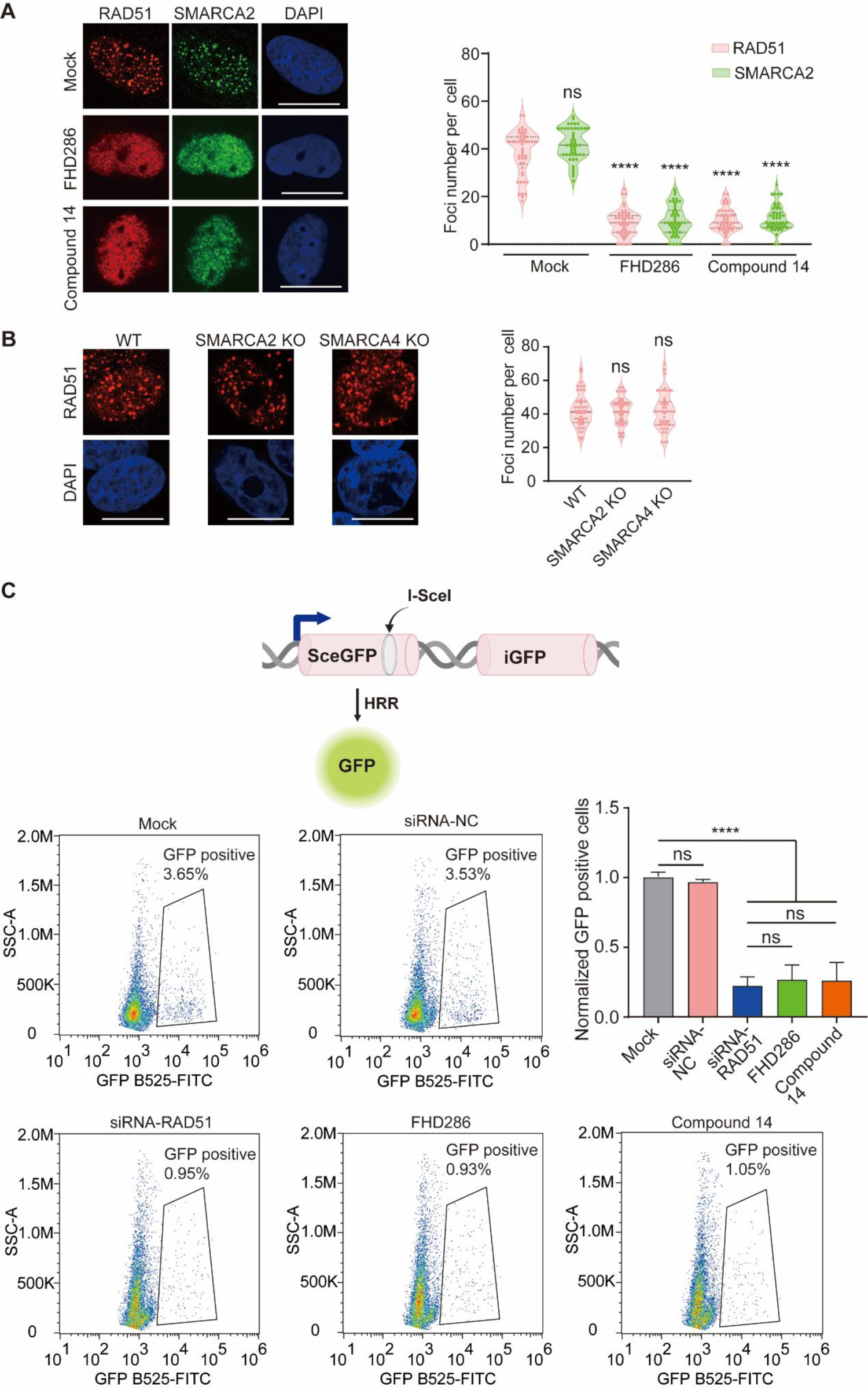
Suppression of the enzymatic activities of SMARCA2/4 impairs RAD51-dependent HR repair. **(A)** HeLa cells were pre-treated with or without FHD286 (1 μM) or Compound 14 (1 μM) for two hours followed by 5 Gy of IR. Foci formation of RAD51, SMARCA2 were examined. **(B)** SMARCA2 or A4 were KO in HeLa cells. IR-induced foci of RAD51 were examined. **(C)** HR efficacy was examined by GFP reporter assays. U2OS cells with DR-GFP insertion were pretreated with or without FHD286 (1 μM) or Compound 14 (1 μM). I-SceI were expressed to induce DSBs. GFP positive populations were calculated by flowcytometry.

**Figure 5.**
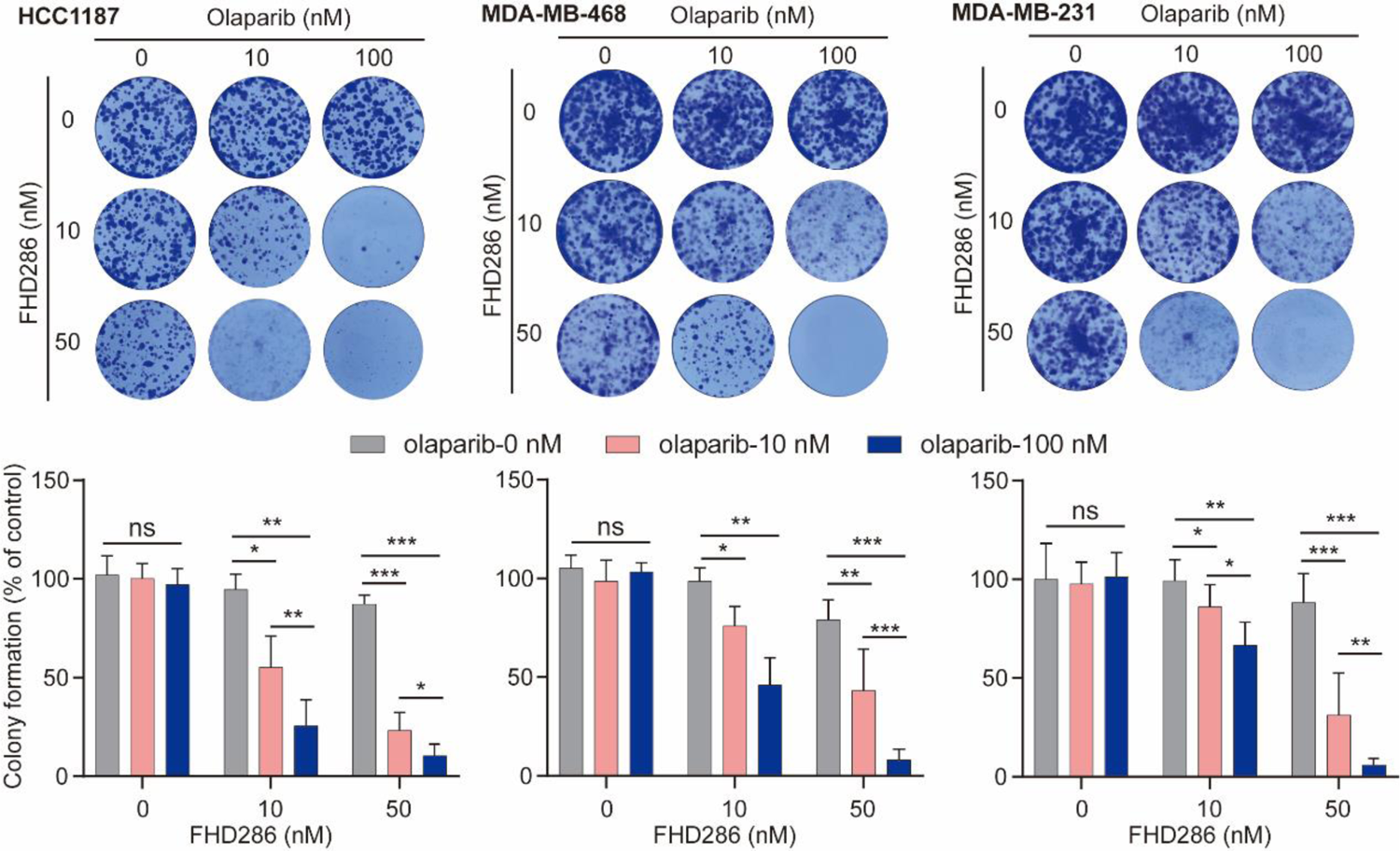
FHD286 sensitizes PARPi for the suppression of tumor cell growth. 1000 cells were seeded in each well, and treated with indicated doses of FHD286 and/or olaparib. Colony formation was examined at Day 14 with 0.1% crystal violet staining. Statistical analyses were shown in the lower panel.

HR-deficient cells are often hypersensitive to PARP inhibitor treatment. To date, PARP inhibitors have been utilized for the treatment of breast, ovarian, pancreatic and prostate cancers with HR deficiency. Interestingly, FHD286 is currently in clinical trials for cancer treatment. Since FHD286 treatment impairs HR repair, we ask if FHD286 treatment can sensitive tumor cells to PARP inhibitor treatment. We selected three breast cancer cell lines (HCC1187, MDA-MB-468 and MDA-MB-231) that are not sensitive to PARP inhibitor olaparib. Interestingly, with 50 nM of FHD286, all these cells line were sensitive to 100 nM olaparib treatment, indicating that FHD286 may have additive effect with PARP inhibitors for cancer treatment in the future.

## Discussion

In this study, we have demonstrated that the enzymatic activity of SMARCA2/4 plays a key role for the recruitment of the SWI/SNF complex to DNA lesions to facilitate DSB repair. Earlier studies have shown other factors such as PARylation, acetylation or phosphorylation events mediate the recruitment of the SWI/SNF complex(16–19). Here, we have shown that only the ATPase activities of SMARCA2/4 are essential for the recruitment of this chromatin remodeling complex. Of note, chromatin remodeling should be one of the early DNA damage response events. Only after the removal of the nucleosome barrier, can the repair machinery access DNA lesions for repair(8,25). Since the ATPase domains of SMARCA2/4 directly interact with genomic DNA(10,13), it is possible that the SWI/SNF complex constantly scans on genomic DNA, and is able to detect DSBs. It will also allow quick chromatin remodeling for damage repair in response to DSBs. Using the laser microirradiation, we found that the relocation of SMARCA2/4 as early as around one second, further demonstrating that SMARCA2/4 are one of earliest DNA damage response factors. Thus, we favor the possibility that SMARCA2/4 are able to quickly and directly recognize DNA lesions as DNA damage sensors.

Earlier studies have shown that lacking SMARCA2 is sufficient to impair HR repair(5,6). However, numerous analyses have suggested that SMARCA2 and SMARCA4 are paralogs and have redundant functions in the SMARCA2/4(2,3,11,22). In our study, we have to abolish both enzymatic activities of SMARCA2/4 to suppress HR repair. Of note, *SMARCA4* gene is often deleted during tumorigenesis, and loss of SMARCA4 is wildly existing in varieties of tumor cell lines(21,26). Thus, if the cells only contain SMARCA2, ablating SMARCA2 is sufficient to disrupt the function of the SWI/SNF complex and impair HR repair. Thus, our results may not be contradicted with earlier studies. Instead, we did more careful and comprehensive analyses by including the consideration of the redundant function of SMARCA4.

In our study, we have shown that suppression of SMARCA2/4 impairs the foci formation of several upstream DNA damage response factors, such as γH2AX, RNF8 and BRCA1, but abolishes the foci formation of downstream repair machinery, such as RAD51. In response to DSBs, the biological function of the SWI/SNF complex is to remove nucleosome barrier and expose naked DNA for DNA end processing(13,15). Usually, at DSB ends, a few hundred base pairs are processed into ssDNA and coated with RAD51 as RAD51-ssDNA filament for strand invasion(27,28). Thus, it is estimated that only a few nucleosomes at DSB ends are evicted by chromatin remodeling complexes. Thus, without enzymatic activities of SMARCA2/4, we could not detect RAD51 at DSBs. However, different from bona fide repair machinery, γH2AX mediates DNA damage response signals extended up to megabases away from DSBs(29–32). Thus, the enrichment of γH2AX is not controlled by the SWI/SNF complex. Instead, because of delayed DSB repair by suppression of SMARCA2/4, these response factors stay at DSBs for prolonged time. Collectively, our studies provide an in-depth understanding on the biological functions of SMARCA2/4 in the context of DNA damage repair.

## Material and Methods

### Plasmids

Human full-length SMARCA2, SMARCA4, SMARACC1 or SMARCD1 sequence was obtained from NCBI (Gene ID: NM_003070, NM_001128849, NM_003074, or NM_003076, respectively) and synthesized by Tsingke Biotechnology company. The full-length SMARCA2 or SMARCA4 cDNA or the fragments of SMARCA2 (N-Ter, amino acids 1-653; Hel, amino acids 654-1383; BRD, amino acids 1384-1590) were cloned into EGFP vector for the expression in mammalian cells.

### Cell culture and transfection

HEK293T and HeLa were cultured at 37 °C with 5% CO_2_ in DMEM supplemented with 10% fetal bovine serum and 1% penicillin/streptomycin. EGFP constructs transfection were performed using Lipofectamine® 2000 (Invitrogen). For SMARCA2 or SMARCA4 knockout, the gRNA sequence of the SMARCA2: 5’-TCCCATCCTATGCCGACGAT-3’, the gRNA sequence of the SMARCA4: 5’-CCGGCGAGGGACCCGGGCTA-3’. For RAD51 gene knockdown, the sequence of si-RAD51: 5’-CUAAUCAGGUGGUAGCUC AUU-3’. The gRNAs and small interfering RNA (si-RNA) were synthesized by Tsingke Biotechnology company. Si-RNA transfections were carried out using Lipofectamine RNAiMAX (Invitrogen), according to the manufacturer’s instructions.

### Antibodies

The following antibodies were purchased from respective companies: anti-γH2AX (Novus, NB100-384), anti-RAD51 (Invitrogen, MA1-23271), anti-RNF8 (14112-1-AP, Proteintech), anti-BRCA1 (NB100-404, Novus), anti-SMARCA2 (Sigma, HPA029981).

### Laser microirradiation assays

GFP-tagged constructs were transfected into the indicated cells which were plated on 35-mm glass bottom dishes. Cellular DNA damages were generated according to the previous study(33). For inhibitor treatment, cells were treated with different inhibitors at 1 μ M for 2 hours followed by laser microirradiation.

SMARCA2/4 inhibitor (FHD286, HY-144835), ATM inhibitor (KU55933, HY-12016), ATR inhibitor (VE-821, HY-14731), CBP/p300 inhibitor (CBP/p300-IN-21, HY-155229) and PARP1/2 inhibitor (Olaparib, HY-10162), were purchased from MedChemExpress. The green fluorescence strips were recorded and then analyzed with ImageJ software. All results represented images of 20 cells from three independent experiments.

### Immunofluorescence (IF) staining

Cells growing on glass coverslips were irradiated with 5 Gy and recovered for 12 h before immunofluorescence. Then, cells were treated with fixation buffer (4% PFA, 0.1% Triton X-100). The slides were incubated with different primary antibodies in 1% BSA at room temperature for 2 h or 4 °C overnight. Subsequently, the slides were washed with PBS and incubated with secondary antibody. DNA was stained and coverslips were mounted with DAPI Fluoromount-G™ (Yeason, 36308ES20). For inhibitor treatment, cells were treated with different inhibitors at 1 μM for 2 hours followed by ionizing irradiation. Compound 14 were synthesized according to the compound structure(34).

### HR efficiency assays

The Tet-DR-GFP-U2OS cells were used which has been reported in the previous study(32). The cells were transfected with pCW-HA-I-SceI-GR plasmid and 1 μg/ml doxycycline was added to the medium. Triamcinolone acetonide (TA) applied to help enzyme cut the I-SceI sites. HR efficiency were quantified by flow cytometry.

### Clonogenic assays

For clonogenic cell proliferation assay, cells were seeded in 6-well plate (1000 cells/well). Then cells were treated with vehicle control or drugs with indicate concentration on the next day and culture medium was refreshed every 3 days for 14 days in total. At the endpoints of colony formation assays, colonies were washed with PBS and fixed with 70% methanol for 10 min, followed by staining with 0.1% crystal violet. The number of colonies was counted by ImageJ software, and survival graphs were generated from three replicate wells with colony numbers normalized to untreated controls.

### Statistical analysis

Statistical analyses were performed using GraphPad Prism7. The statistical significance carried out by two-tailed unpaired Student’s t-test. All of the data were presented as mean ±SD. The significance levels are annotated: *, P≤0.05. **, P≤0.01. ***, P≤0.001. ****, P≤0.0001.

### Author contributions

L.Y. and D.W. designed the project, performed the experiments, analyzed data and wrote the manuscript. Both authors have read and approved the content of the submitted manuscript.

## Data availability

No data was used for the research described in the article.

## Acknowledgement

This work was supported in part by grants from Hangzhou City Leading Innovation and Entrepreneurship Team (TD2020004), “Pioneer” and “Leading Goose” R&D Program of Zhejiang (2024SSYS0033), Westlake University Education Foundation, and Westlake Laboratory of Life Sciences and Biomedicine. We apologize to all authors whose work could not be cited due to space limitations. Figures were created with BioRender.com.

