## Supplemental Figure for "SMARCA2 and SMARCA4 participate in DNA damage repair"

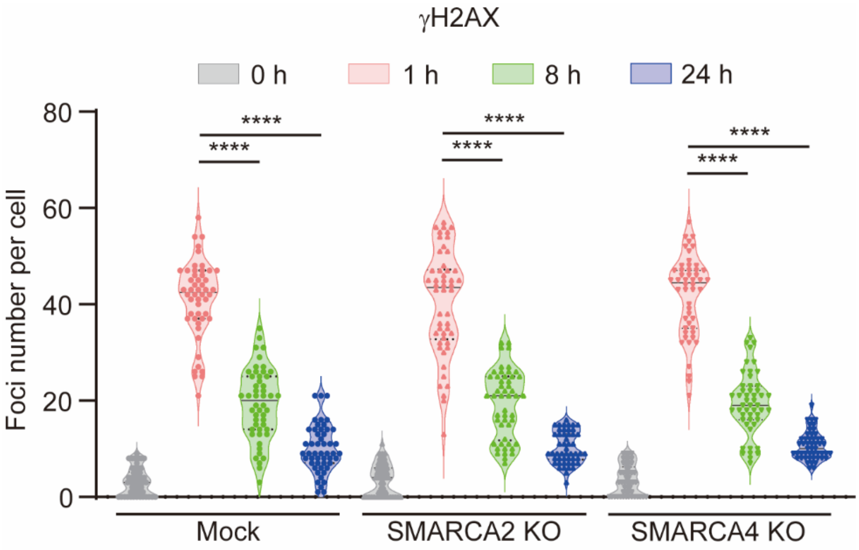


**Figure S1. Lacking either SMARCA2 or SMARCA4 does not affect the foci formation of γH2AX.**

Following 5Gy of IR treatment, the foci of γH2AX were examined at the indicated recovery time points.
